# Laser-integrated nanophotonic neural probes with on-chip sensors for addressable photostimulation

**DOI:** 10.64898/2026.07.24.740534

**Authors:** David A. Roszko, John N. Straguzzi, Homeira Moradi Chameh, Mayra Santos da Silva, Piyush Kumar, Xin Mu, Hongyao Chua, Robert Lawrowski, Frank Weiss, Guo-Qiang Lo, Mariel Jama, Joyce K. S. Poon, Taufik A. Valiante, Wesley D. Sacher

## Abstract

Studying the role of individual neurons in behavior and disease requires tools for controlling neural activity with high spatiotemporal resolution. Implantable nanophotonic neural probes are capable of delivering targeted photostimulation to enable genetically distinct neurons to be selectively controlled, yet face barriers to achieving scalable emitter densities and lack sensors for monitoring feedback signals relevant to device operation. To address this, we developed laser-integrated nanophotonic neural probes, which feature hybrid-integrated laser diodes (LD) and thermo-optic photonics switches for scalable emitter addressing and on-chip sensors for monitoring optical power and temperature during photo-stimulation. Devices were fabricated at a commercial silicon photonics foundry on 200-mm diameter silicon-on-insulator (SOI) wafers in an active visible-light platform and were controlled using a custom-developed electronic circuit board. Each device features a hybrid-integrated InGaN LD which couples 450-nm light into a reconfigurable photonic switching tree for delivering spatially resolved photostimulation through 16 emitters along a 3-mm implantable shank. Using the on-chip photodetectors, we demonstrate how devices can enable switching tree calibration as well as output power monitoring during photostimulation. Furthermore, using the on-chip temperature sensors, we show how device temperature perturbations resulting from LD and thermo-optic switch activation can be directly monitored during photostimulation to ensure temperature fluctuations remain below 1 **°**C. We validate our design by delivering high spatiotemporal photostimulation during an *in vivo* optogenetic experiment with simultaneous Neuropixels recording to monitor evoked responses. Overall, these scalable integrated devices offer a pathway for neuroscientists to conduct fiberless optogenetic experiments with greater control and precision.

## 1 Introduction

Optogenetics is a powerful technique in neuroscience which enables genetically distinct neurons to be controlled and studied using light ^1^. To facilitate optogenetic experiments in deep brain regions (depth ≥ 1 mm), researchers are developing a myriad of implantable tools for delivering light into the brain. These implantable devices use microscale light-emitting diodes (µLEDs or OLEDs) ^2–5^, optical fibers ^6–11^, or photonic waveguides ^12–18^ to directly emit light or guide light into the brain to deliver targeted photostimulation. Each device architecture presents clear trade-offs and designing devices with high optical power, spatiotemporal resolution, and small cross-sectional areas remains challenging.

Nanophotonic neural probes are planar, implantable devices which route light into the brain using integrated nanophotonic waveguides ^13,14,16–19^. Compared to other implantable devices, nanophotonic neural probes offer several key advantages for optogenetic experiments. Like optical fibers, the broad optical transparency of dielectric waveguide materials, most often silicon nitride (SiN), allows nanophotonic devices to deliver targeted photostimulation across the visible spectrum at optical powers sufficient for driving local and regional neural activity ^16–18^. Yet unlike optical fibers, photonic neural probes also benefit from the dense circuit integration afforded by modern electronic and photonic foundry processes. By routing light with photo-lithographically defined waveguides with high optical confinement, nanophotonic neural probes can implement dense arrays of light emitters while maintaining small cross-sectional areas. Moreover, because photonic neural probes can be fabricated with complementary metal-oxide semiconductor (CMOS) compatible processes, devices can be co-integrated with active electronics for on-chip modulation and sensing ^20,21^ and dense active electrode arrays for high resolution electrophysiological recording in addition to multichannel photostimulation ^15,18^. Furthermore, by leveraging the capabilities of foundry fabrication processes, nanophotonic neural probes benefit from enhanced scalability, supporting continuous improvements in circuit complexity and improved emitter densities while also offering a pathway to scalable manufacturing volumes to support broad distribution to the neuroscience community. Early demonstrations of nanophotonic neural probes have relied on fiber-coupled external laser systems and scanning optics and/or wavelength tuning to deliver optical power and to select optical channels ^12,14,16,17,19,22^. Although these external systems provide precise optical control, they also increase the complexity of probe operation, alignment, packaging, and maintenance. This complexity can limit adoption by neuroscientists and reduce the practical impact of otherwise powerful nanophotonic probe technologies. Integrated photonics offers a path to improve usability by transferring much of this system-level complexity from external instrumentation onto the probe itself. Recent state-of-the-art nanophotonic devices have begun to do this by using integrated thermo-optic photonic switching trees to reconfigurably address multiple on-chip light emitters, thereby eliminating the need for scanning optics for channel addressing ^13,15,18^. Because each additional switching layer can increase the number of addressable outputs exponentially, photonic switching trees provide a scalable approach for high density emitter integration, within the limits of propagation losses and waveguide crosstalk. However, devices featuring photonic switching trees have still relied on fiber-coupled external laser sources for light delivery, complicating device packaging and limiting scalability. Hybrid laser integration has been explored as an alternative to optical fiber coupling in photonic neural probes ^23,24^, but past implementations required one LD per output channel. This architecture also can create significant thermal constraints, with thermal simulations indicating excessive heating during extended operation, necessitating strict limits on laser duty cycle and amplitude to avoid thermal effects on neural tissue ^24^. In addition to integrated optical sources and switches, practical implantable photonic devices would benefit from on-chip sensors for monitoring optical power and device temperature. Optical power sensors would improve device control and usability by enabling calibration and output power stabilization, while temperature sensors would allow users to directly monitor heating from active circuit elements during experiments which could affect neural responses ^25,26^. To date, however, sensor integration in nanophotonic neural probes has only been reported preliminarily for devices with ^27^ and without ^28^ integrated LDs. Together, integrated switching, integrated optical sources, and on-chip sensing would move key functionalities from bulky external systems onto the neural probe itself. Combining these capabilities in a single integrated device would therefore enable highly scalable, fiberless nanophotonic neural probes that are easier to package, operate, calibrate, and use in neuroscience experiments.

Toward this aim, we present laser-integrated nanophotonic neural probes: photonic microsystems featuring hybrid integrated lasers and thermo-optic photonic switches for reconfigurable emitter addressing, as well as on-chip optical power and temperature sensors for monitoring device performance and heating during targeted photostimulation. By combining both scalable integrated optical source and switching architectures, these devices offer a pathway for mass-manufacturable devices with greater emitter densities for achieving high-resolution photostimulation. Here, we present the development and characterization of a prototype device which we characterize through both bench and *in vivo* experiments. Through coordinated activation and monitoring of the integrated laser, photonic switching tree, and on-chip photodetectors, we show how devices can enable photonic circuit calibration and deliver photostimulation pulse trains with simultaneous output power monitoring. We also show how integrated temperature sensors can be monitored during photostimulation pulse trains to ensure device temperature perturbations remain below 1°C, thus limiting the thermal effects on neural tissue during *in vivo* experiments. We validate our design by delivering targeted photostimulation during an *in vivo* optogenetic experiment with simultaneous Neuropixels ^29^ recordings for measuring evoked neural responses. An initial report of these results was presented in our conference abstract ^27^.

## 2 Results

### 2.1 Laser-integrated nanophotonic neural probe design and system architecture

The prototype nanophotonic neural probes with integrated lasers, photonic switches, and on-chip sensors were realized in our active visible light silicon photonics platform ^20,21,30^ (Fig. 1(a)). Each neural probe photonic integrated circuit (PIC) included a hybrid integrated laser diode (LD) coupled to a photonic switching tree for programmable channel selection (Fig. 1(b)). Additionally, each device featured on-chip photodetectors for calibration and monitoring of the photonic circuitry as well as thermistors for measuring device heating during operation. Using these functionalities, the neural probes were programmed to deliver addressable photostimulation through discrete grating coupler emitters for spatially resolved photostimulation (Fig. 1(c)). When paired with stand-alone devices for high-resolution electrophysiological recording ^29^, the laser-integrated nanophotonic probes enable simultaneous targeted photostimulation, optical power monitoring, and temperature sensing to support *in vivo* experiments in optogenetic mice (Fig. 1(d)).

**Fig. 1.**
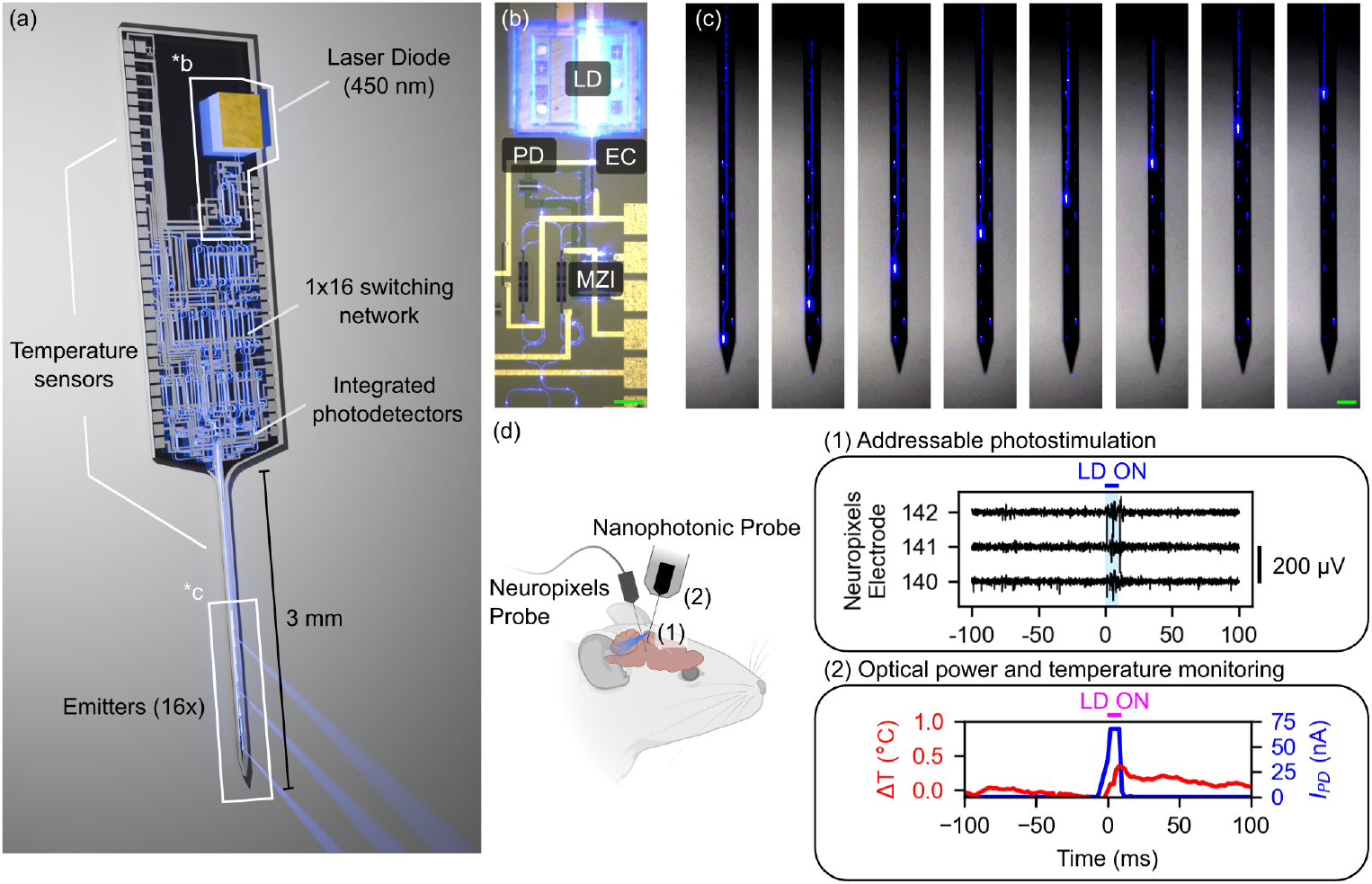
Overview of the laser-integrated nanophotonic neural probe design. (a) Conceptual rendered image of the laser-integrated nanophotonic neural probe with labeled components. (b) Micrograph of hybrid-integrated InGaN laser diode (LD) on a neural probe. Light from the LD is coupled into the photonic integrated circuit via an edge coupler (EC), where it is detected by an evanescently-coupled photodetector (PD) and directed using a Mach-Zehnder Interferometer (MZI) thermo-optic photonic switch. Scale: 100 µm. (c) Micrograph of reconfigurable emitter selection at the shank of the device. Scale: 100 µm. (d) Conceptual demonstration of device operation during dual-probe experiment. The nanophotonic probe monitors device temperature and optical power during targeted photostimulation while photostimulation-evoked activity is measured simultaneously using a Neuropixels 1.0 probe ^29^. Components of this figure were created with BioRender.com.

The devices were fabricated on 200-mm diameter silicon-on-insulator (SOI) wafers at Advanced Micro Foundry (Singapore), which featured SiN waveguides for photonic routing and active circuitry for modulation and sensing (Fig. 2(a), see Sec. 4.1 for details). The design featured a blue (450 nm) hybrid integrated InGaN laser diode (LD) ^30,31^ coupled to a 1x16 photonic switching tree, implemented with Mach-Zehnder interferometer (MZI) switches with thermo-optic phase shifters, for reconfigurable channel selection (Fig. 2(b)). In addition to the primary LD input, light could also be coupled into the photonic switching tree using a secondary edge coupler input for characterization purposes. Following methods outlined in ref. 30, LDs were integrated onto each PIC using a passive-alignment flip-chip bonding process into customized LD trenches, featuring SOI mechanical stoppers for precision height alignment, in-plane alignment markers, and under-bump metallization (UBM) with AuSn-alloy solder for making electrical contacts, to achieve high throughput optoelectronic assembly. Building on our previous work, the arms of the thermo-optic photonic switches were designed with multiple loopbacks (3 passes) and lateral thermal isolation trenches to improve thermo-optic power efficiency ^20^. At the output of the photonic switching tree, each of the 16 output branches was routed to a discrete grating coupler emitter on the implantable device shank for spatially-resolved optogenetic stimulation (16 channels arranged in two columns, pitch (lateral): 46 µm, pitch (depth): 90 µm; tapered shank width: 90 µm to 82 µm, shank length: 2.97 mm, thickness: 60 µm). Additionally, each output branch was also coupled to an optical power tap, which routed a small fraction (approximately 5%) of the light to an evanescently-coupled photodetector ^21^ for optical power monitoring. And finally, the design also included doped Si thermistors near the integrated LD and on the device shank for measuring the device temperature during operation. By simultaneously activating the integrated LD and photonic switching tree (Fig. 2(c)), we confirmed that the neural probes could be configured for addressing multiple outputs (Fig. 2(d)) or individual emitters (Fig. 2(e)).

**Fig. 2.**
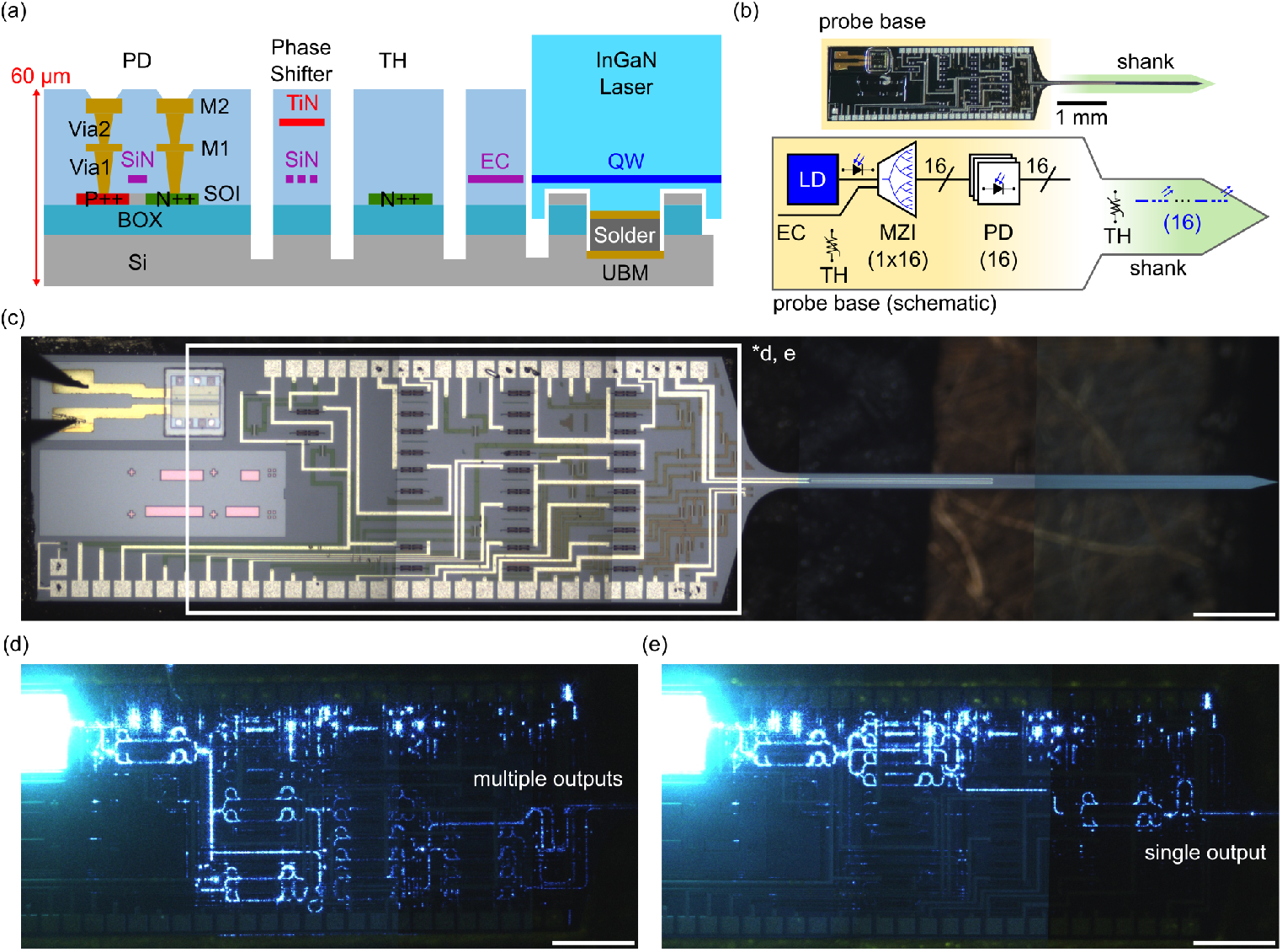
Laser-integrated nanophotonic neural probe design. (a) Cross-sectional diagram of the active visible-light silicon photonic platform used to define the neural probe. (b) Schematic diagram of the neural probe photonic integrated circuit. A hybrid-integrated LD couples light into a 1x16 photonic switching circuit, composed of MZI photonic switches, which routes light to 16 separate grating coupler emitters for targeted photostimulation. Each output branch of the switching tree features an evanescent power tap, which couples ∼5% of the light to an evanescently-coupled PD. Doped silicon thermistors measure device temperature at the base and shank of the probe. (c) Stitched micrograph of laser-integrated nanophotonic neural probe. Scale bar: 500 µm. (d, e) Stitched micrographs of laser and photonic switching tree activation for (d) multiple and (e) single output operation. Scale bars: 500 µm. PD - photodetector, M - aluminum metal, SiN - silicon nitride, BOX - buried oxide, SOI - silicon-on-insulator, TiN - titanium nitride, TH - thermistor, EC - edge coupler, QW - quantum wells, UBM - under-bump metallization, MZI - Mach-Zehnder interferometer photonic switches.

To interface with the integrated nanophotonic neural probes, we also developed a custom control PCB and modular carrier PCBs for each neural probe to control the on-chip lasers and photonic switching tree while monitoring signals from the integrated photodetectors and thermistors (Fig. 3(a)). The control PCB included two 16-channel 16-bit voltage-output digital-to-analog converters (DACs) (DAC81416, Texas Instruments, Dallas, TX, USA) for controlling the 15 on-chip thermo-optic photonic switches, and one 4-channel 10-bit current-output DAC (ISL78365, Renesas Electronics Corp., Tokyo, Japan) driving the LD. Additionally, for device monitoring, the control PCB included one 64-channel 20-bit current-input analog-to-digital-converter (ADC) (DDC264C, Texas Instruments) for monitoring the on-chip photodetectors, and two 32-bit differential voltage-input ADCs in a 3-wire RTD configuration (ADS1263, Texas Instruments) for monitoring the thermistors. Importantly, we designed the control PCB with more LDD, DAC, and ADC channels than strictly required by the current laser-integrated neural probe to ensure compatibility with future probe designs and streamline further developments. A shared serial peripheral interface (SPI) bus, SPI1, was used to read and write data to the two DACs (daisy-chain configuration), the two thermistor ADCs, and for programming the LDD. The photodetector ADC was placed on a dedicated SPI bus (SPI2) to permit continuous photodetector monitoring during device operation. The LDD drive current was controlled by a dedicated 10-bit parallel data interface (PI) to enable seamless laser activation during continuous data monitoring on SPI1. The control board was designed to interface with a commercially available field-programmable gate array (FPGA) development module (CMOD-S7, Digilent co NI, Austin, TX, USA), featuring a Xilinx Spartan-7 FPGA, capable of controlling and monitoring the signals from all on-chip components. Finally, data was sent to and from the control PCB over an additional SPI connection to a computer via a microcontroller. For interfacing with the control PCB, each integrated nanophotonic neural probe was mounted onto a dedicated carrier PCB, which could be interchangeably connected to the control PCB via a high-density interface (HDI) connector (Fig. 3(b,c)). After packaging the neural probes, we then used this system for delivery and monitoring of programmable photostimulation pulse trains.

**Fig. 3.**
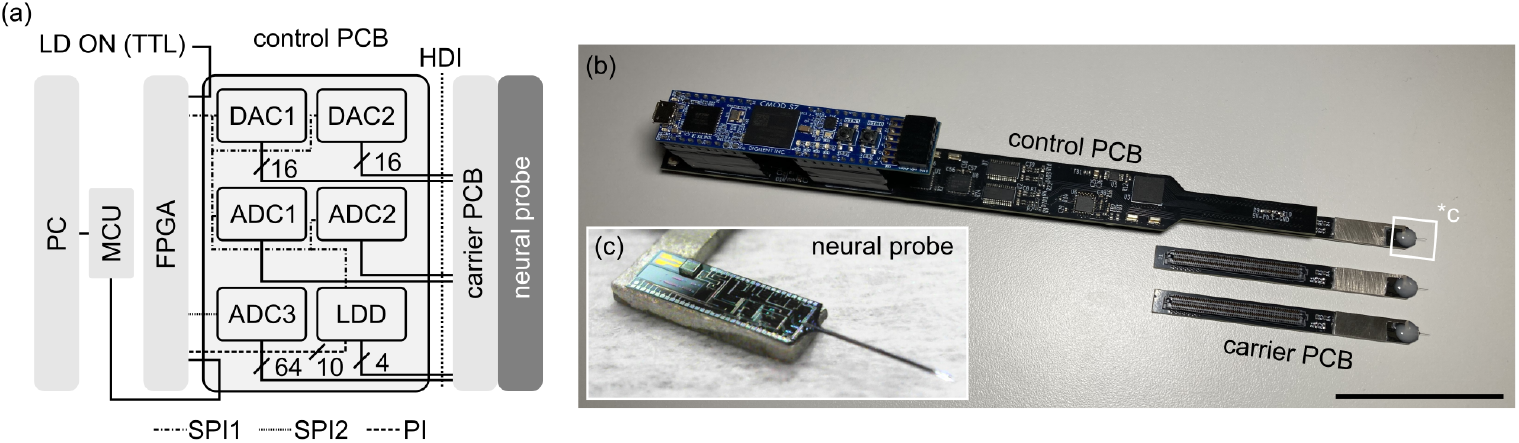
Peripheral electronic hardware for neural probe control and data acquisition. (a) Schematic diagram of the control PCB. (b) Photo of the control and carrier PCBs. Scale: 50 mm. (c) Photo of a neural probe device prior to attachment onto a carrier PCB. PC - computer, MCU - microcontroller, FPGA - field-programmable gate array, DAC - digital-to-analog converter, ADC - analog-to-digital converter, LDD - laser diode driver.

### 2.2 Integrated photodetectors enable photonic switching circuit calibration and optical power monitoring during photostimulation

By using the integrated PDs for optical power monitoring, we demonstrate how these sensors aid in two critical device functions: photonic switching tree calibration, wherein the bias points for each MZI are identified using feedback from the PDs, and optical power monitoring during photostimulation pulse trains. For both of these functions, coordinated activation and monitoring of the integrated LD, photonic switching tree, and PDs was managed by dedicated finite state machines implemented on the control PCB FPGA.

For calibration of the photonic switching tree, a linearly-ramped bias voltage was applied to the first switch in the switching tree (MZI11; layer 1, switch 1) while integrated PDs monitored the optical power at each of the 16 output branches of the switching tree. For these measurements, the LD was operated in continuous wave (CW) mode at a constant, low-amplitude drive current (20-23 mA) to permit constant measurements while reducing device heating. As the thermo-optic phase shifter power increased, light was alternately coupled to each output of MZI11, which was observed as an alternating PD signal for each branch (Fig. 4(a)). Focusing on two representative PD signals from Fig. 4(a) (1 per MZI output), bias values for maximizing light in either of the two MZI outputs were identified by locating the PD signal maxima coupled to each respective branch (Fig. 4(b)). This process was then repeated sequentially for each switch in the tree until all bias points were identified, thus enabling any of the 16 emitters to be reconfigurably selected. Additionally, by measuring the change in MZI power between PD signal maxima at opposite ports, we calculated the power required for switching from one MZI output port to another (P*_π_*, power required for a *π* phase shift in the thermo-optic phase shifter) (Fig. 4(c)). We then extracted the P*_π_* values for each switch and found that the average P*_π_* value was 18 1 mW (n=62 switches from 5 neural probes) (Fig. 4(d)).

**Fig. 4.**
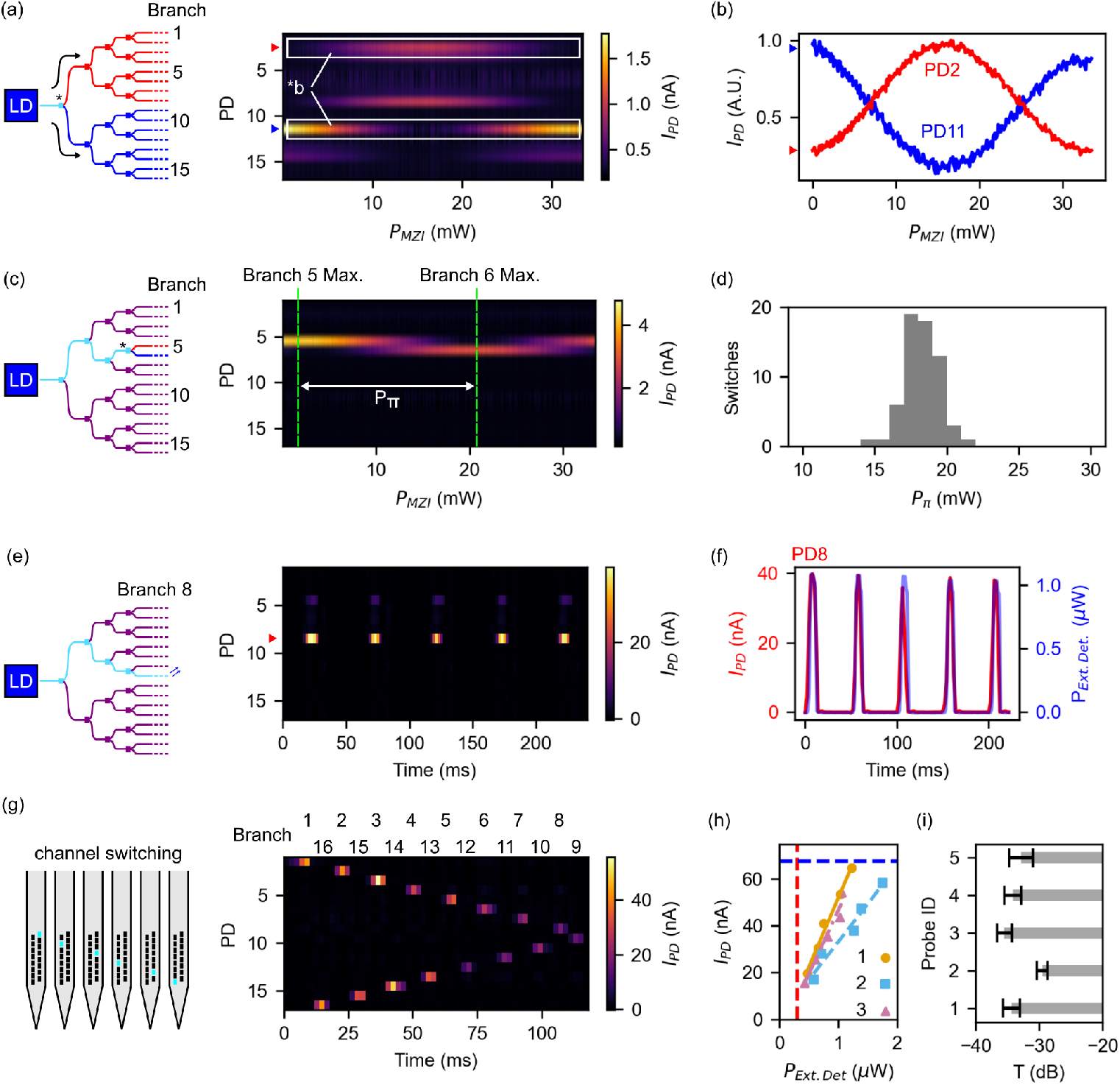
Demonstration of laser diode (LD) and photonic circuit operation with output monitoring via integrated photodetectors (PDs). (a) Calibration measurement for MZI11 (label: *, layer 1, switch 1). Photocurrent across all branch PDs (*I_PD_*) is shown vs. MZI drive power (*P_MZI_* ). *I_P D_* maxima correspond to *P_MZI_* values for optimal routing to next switch layer (for complete switching between illustrated red and blue paths). (b) Representative normalized PD signals from (a). (c) Calibration measurements for MZI43 (label: *, layer 4, switch 3); the MZI switching power, *Pπ* , is labeled. (d) *Pπ* values extracted from calibration data (n=62 switches from 5 devices). (e) Photostimulation pulse train delivered from a calibrated device and monitored using on-chip PDs. (f) Comparison of on-chip and external optical power meter (*P_Ext.Det_*) signals during photostimulation pulse train. (g) Multichannel photostimulation pulse train delivered and monitored using on-chip PDs. (h) Peak integrated on-chip PD photocurrent vs. emitter output power detected with external optical power meter (data shown for n=3 emitters from 1 probe). (i) Average neural probe transmission values (n=56 emitters from 5 neural probes).

After calibration, we then used the integrated photodetectors for output power monitoring during photostimulation pulse trains. First, to determine the maximum light input to the integrated nanophotonic neural probes, we characterized the performance of individual InGaN LDs (n=3) mounted on thermally-conductive submounts (Alpes Lasers, St-Blaise, Switzerland). Light-current-voltage (L-I-V) measurements were made using a source measure unit (SMU) (B2912A, Keysight, Santa Rosa, CA, USA) in response to current pulses (1 ms duration, 50 ms interpulse interval) with linearly increasing amplitude (Fig. S1(a)). Based on these measurements, the threshold current was measured as 10.6±0.4 mA and LDs were observed to output 44±4 mW when driven at 80 mA. After characterizing the LDs, we then measured the thermo-optic switch rise and fall times to estimate the maximum switching frequency permitted by the photonic switching tree. 450 nm light was coupled into the device using an optical fiber while a thermo-optic switch was driven to maximize light in either MZI outputs. A multimode fiber, coupled to a low-rise-time external photodetector (12 MHz bandwidth, PDA36A2, Thorlabs, Newton, NJ, USA), was positioned over a grating coupler emitter connected to the thermo-optic switch to monitor the temporal dynamics of the switch (Fig. S1(b, c)). Following these measurements, an exponential function was fitted to the photodetector data, and the rise and fall times of the photonic switch were estimated from the fitted functions as 96 µs and 59 µs, respectively.

Following these initial measurements, we then programmed a calibrated laser- integrated nanophotonic neural probe to selectively deliver a pulse train (5 pulses, pulse width: 10 ms, pulse period: 50 ms; LD current: 26 mA) through emitter 8 (Fig. 4(e)) while all PDs were sampled every 2 ms. During the pulse train, we observed a highly localized current signal corresponding to the delivered laser pulse train, with a peak current of 40 nA. To determine the relationship between the PD current and the output power at the emitter, we also simultaneously measured the emitter output power directly using an external optical power meter (PM103A, Thorlabs). Based on these paired measurements, we found that the peak PD current corresponded to an output power of 1.1 µW (Fig. 4(f)).

Next, we explored how coordinated activation of the LD, photonic switching tree, and photodetectors could be used to deliver and monitor rapid switching across multiple emitters for delivering targeted spatiotemporal photostimulation. A nanophotonic neural probe was configured to deliver pulses (pulse width: 5 ms, interpulse interval: 1 ms; LD current: 26 mA) sequentially from each emitter with increasing depth (Fig. 4(g)). For this pulse train, we observed highly localized PD activation for each output channel in sequential order, confirming that the device could both deliver and monitor rapid and complex photostimulation patterns.

We then investigated the measurement range of the integrated photodetectors for quantifying the emitter output powers. Similar to the measurements in Fig. 4(f), a calibrated neural probe was programmed to deliver optical pulses with increasing output power (LD current: 24-26 mA) while both the on-chip photodetectors and an external optical power meter, placed directly above the device shank, were monitored. For each photostimulation pulse, we then plotted the peak on-chip photodetector current vs. the measured emitter output power (data for n=3 representative emitters shown in Fig. 4(h)). For the photodetector ADC, given the current firmware implementation (integration time: 2.2 ms, maximum integrated charge per channel: 150 pC), it was known that maximum detectable current was 67.6 nA per channel (indicated with a blue dashed line in Fig. 4(h)). Similarly, for the external optical power meter, we estimated the minimum detectable signal as 0.3 µW, which was calculated as 2× the average standard deviation of the signal (indicated with a red dashed line in Fig. 4(h)). By assessing pulses within this range (n=72), we found that the on-chip PDs could estimate emitter powers up to approximately 1.9 µW before saturating (Fitted slopes in Fig. 4(h): (1) m=59.1; b=-7.1, (2) m=32.5; b=0.7, (3) m=54.6; b=-7.9). While this measurable upper-bound is below the maximum emission power of the probe, calibration of the photonic switching circuit can be performed at lower input laser powers within the range of the on-chip PDs.

Lastly, by comparing the measured emitter output powers to the estimated LD power (see Fig. S1(a)) for several neural probes (n=56 emitters from 5 neural probes), we calculated the average transmission of the neural probes as -33.1±2.4 dB (shown per probe in Fig. 4(i); transmission values: -34.3±1.3 dB, -29.5±0.8 dB, -35.5±1.2 dB, -34.1±1.3 dB, -32.8±1.9 dB), corresponding to an average output power of 22 µW at LD currents of 80 mA. We then analyzed the simulated beam profiles for the grating coupler emitters (Fig. S2) and observed that each emitter produced a narrow, low-divergence beam with an emission angle of 18° relative to normal (Fig. S2(a,b)). Using the beam propagation method in scattering media ^32^, we then estimated the beam dimensions in the top- and side-profiles after propagating through simulated neural tissue (Fig. S2(c-f)), and found that the full-width at half-maximum (FWHM) beam widths after propagating a distance of 100 µm were 7.5 µm (top-profile, Fig. S2(c, e)) and 8.9 µm (side-profile, Fig. S2(d, f)), respectively. By considering the beam as an astigmatic Gaussian, we then estimated the average beam intensity at a distance 100 µm away from the emitter as 144 mW/mm^2^ (estimated beam waist in transverse directions: 6.4 µm, 7.6 µm; estimated Gaussian beam area: 152.8 µm^2^; emitter optical power: 22 µW; LD current: 80 mA).

### 2.3 Integrated thermistors for temperature monitoring of active photonic circuitry during bench and *in vivo* experiments

Using the integrated thermistors, we then investigated the effects of LD and thermo-optic photonic switching tree activation on device temperature. We demonstrate read-out of on-chip temperature sensors during both bench and *in vivo* experiments, confirming that device shank temperatures remain below 1°C when delivering brief photostimulation pulses.

First, we calibrated the shank thermistors for 3 nanophotonic neural probes by immersing the device shanks in a heated water bath relative to a commercial temperature sensor (see Sec. 4.2) (Fig. 5(a)). Based on these calibration measurements, the response of each thermistor was observed to be approximately linear over the measurement range, with an average slope of 5.10e+05 [ADC code*/*°C] (minimum slope: 4.28e+05 [ADC code*/*°C], maximum slope: 6.25e+05 [ADC code*/*°C]) and intercept values specific to each probe. Because the base thermistors had similar nominal impedance values (37.0±0.6 kΩ, n=3) as the shank thermistors (36.2±0.2 kΩ, n=3), we also used these calibration measurements to calculate the relative temperature change at the base thermistor.

**Fig. 5.**
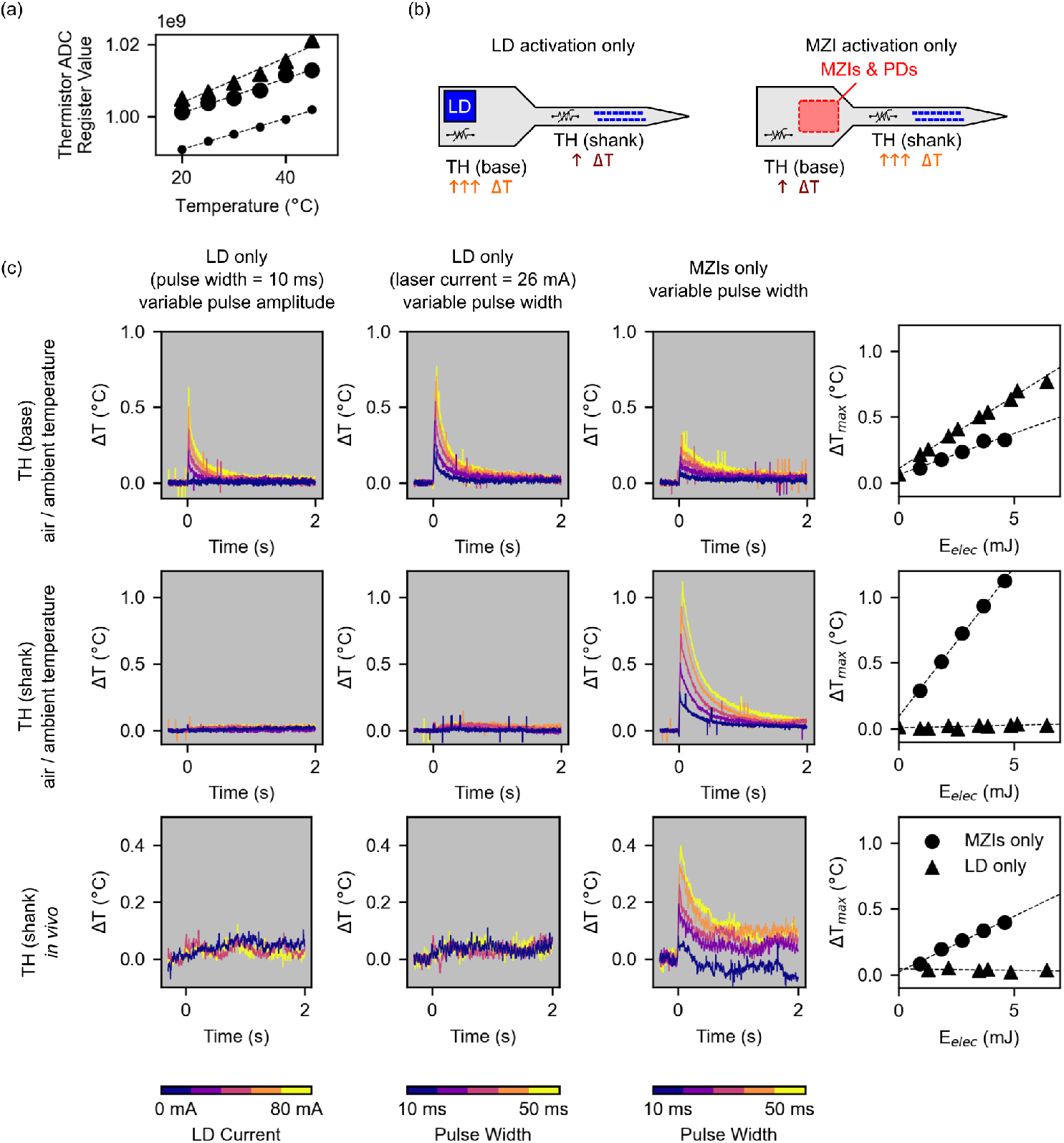
On-chip thermistor (TH) measurements of temperature in response to isolated LD or photonic switching tree activation. (a) Thermistor calibration measurements with probe immersed in a temperature-controlled water bath (n=3). (b) Conceptual illustrations of isolated LD or photonic switching tree activation to assess isolated contributions to device heating. (c) Temperature perturbations at the device base (row 1) and device shank (rows 2 and 3) in response to pulsed operation of the LD (column 1: 10 ms pulse width, variable amplitude; column 2: LD current fixed at 26 mA, variable pulse width) or photonic switching tree (column 3: 4 MZIs activated for selecting branch 1; variable pulse width). Peak temperature perturbation vs. electrical energy delivered to the probe within each pulse (E*_elec_*) is shown in column 4. Rows 1 and 2: probe in ambient air. Row 3: probe implanted *in vivo* (as in Fig. 7).

Following thermistor calibration, we assessed the contribution of device heating caused by activating either the LD or photonic switching tree in isolation while the device was in ambient air (Fig. 5(b, c)). Focusing first on the base thermistor, which was closest to the hybrid integrated LD, we programmed an integrated nanophotonic neural probe to emit single laser pulses with variable pulse amplitude (LD current: 0 mA – 80 mA, pulse width: 10 ms) or variable pulse width (pulse width: 10 ms – 50 ms, LD current: 26 mA) while monitoring the response of the thermistor every 2 ms (Fig. 5(c), row 1). In both cases, LD activation resulted in a sharp increase in the base temperature, which increased in magnitude with increasing electrical energy consumption of the LD (Fig. 5(c), row 1, columns 1 and 2). We then activated the photonic switching tree in isolation while the LD was switched off, to investigate the contribution of device heating from the thermo-optic switches. For these experiments, the switching tree was activated in the configuration for selecting channel 1 (pulse width: 10 ms – 50 ms), which was calculated to dissipate the most electrical power of any switching configuration for this particular probe (92 mW), while the base thermistor was monitored every 2 ms (Fig. 5(c), row 1, column 3). Similar to the LD activation, this resulted in rapid heating at the base of the device, although less prominently than the LD activation. For comparison, we then calculated the peak temperature excursion as a function of the energy consumption of each pulse (Fig. 5(c), row 1, column 4), and observed that the temperature of the base of the device was affected more prominently by LD activation (fitted slope: 0.11 °C*/*mJ) compared to the photonic switching tree activation (fitted slope: 0.06 °C*/*mJ).

After assessing device heating at the base of the device, we then investigated the thermal effects of isolated LD and photonic switching tree activation on the shank of the device, which would be directly in contact with neural tissue during *in vivo* experiments (Fig. 5(c), row 2). Using the same pulse conditions described above, we programmed an integrated nanophotonic neural probe to undergo pulsed activation of either the LD (Fig. 5(c), row 2, columns 1 and 2) or the photonic switching tree (Fig. 5(c), row 2, column 3) while the shank thermistor was sampled every 2 ms. Notably, we observed minimal heating at the device shank during laser activation, with peak temperature excursions below 0.1 °C (Fig. 5(c), row 2, columns 1 and 2). Because the peak temperature excursion was less distinct for these measurements, the maximum temperature excursion was calculated as the average temperature value between 1 - 2 seconds after the pulse. In contrast, activating the photonic switching tree resulted in rapid heating at the device shank, with peak temperature excursions exceeding 1°C as pulse widths extended beyond 50 ms (Fig. 5(c), row 2, column 3). Comparing the peak temperature excursions for both the LD and photonic switching tree as a function of energy consumption, we observed that the shank temperature rose rapidly when energy was consumed by the photonic switching tree (fitted slope: 0.23 °C*/*mJ) but not when the LD was activated (fitted slope: *<*0.01 °C*/*mJ) (Fig. 5(c), row 2, column 4).

Following these experimental measurements, we investigated in simulation (COMSOL Multiphysics, COMSOL AB, Stockholm, Sweden) how to reduce the temperature increase in the probe shank. The neural probe was simulated and transient heating was assessed in response to activation of the LD (Fig. S3(a)) or photonic switching tree (Fig. S3(b)). For these simulations, point measurements were recorded at the base and shank thermistors as well as at points associated with metal wiring connections which might shunt heat directly to the thermistors (Fig. S3(c,d)). For our simplified model, which did not directly simulate the intricate metal wiring of the photonic chip, we observed that matched experimental data agreed well with the simulated results, which exhibited features of both rapid heating via the metal shunt pathways and gradual heating at the thermistors themselves (Fig. S3(e-h)). Similar to the experimental data, we observed rapid heating at the base thermistor in response to LD activation, with reduced heating at the device shank, and more pronounced heating from the photonic switching tree at the device shank compared to the probe base (Fig. S3(i,j)). Focusing on the probe shank, we then compared the effects of device heating for the current probe design and an extended design with slightly longer base (1 mm longer), which would have the added benefits of a larger base area for improved heat sinking as well as moving the active circuit components further away from the shank (Fig. S4(a,b)). Here, we observed that the extended base design lowered the peak shank temperature perturbation from MZI activation from 0.28 °C to 0.08 °C (Fig. S4(c)), and similarly lowered the peak shank temperature perturbation from LD activation from 0.12 °C to 0.08 °C (Fig. S4(d)).

Next, we investigated the thermal effects of isolated activation of the LD or photonic switching tree on the shank temperature with the device implanted *in vivo*. Following methods outlined in Sec. 4.3, a nanophotonic neural probe was implanted in cortex of an awake, head-fixed, Thy1-ChR2^33^ mouse while a series of pulses were delivered and the shank thermistor temperature was monitored (Fig. 5(c), row 3; LD only, variable LD current: 0 mA – 80 mA, pulse width: 10 ms; LD only, variable pulse width: pulse width: 10 ms – 50 ms, LD current: 26 mA; photonic switching tree only, pulse width: 10 ms – 50 ms; 3 pulses per setting). Similar to the shank thermistor measurements captured in air (Fig. 5(c), row 2), we observed minimal heating effects from directly activating the LD (Fig. 5(c), row 3, columns 1 and 2; fitted slope in column 4: *<*0.01 °C*/*mJ) and rapid heating when the switching tree was activated (Fig. 5(c), row 3, column 3; fitted slope in column 4: 0.08 °C*/*mJ).

After assessing the unique contributions to device heating from the LD and photonic switching tree alone, we then analyzed the combined thermal effects when both the LD and switching tree were activated, with pulse parameters relevant for optogenetic stimulation (LD current: 80 mA, pulse width: 10 ms) (Fig. 6(a)). In the same awake, head-fixed mouse, a series of photostimulation pulses (n=135 pulses across all 16 emitters) were delivered *in vivo* while the change in temperature at the device shank was monitored. Across all pulses, we observed that the peak temperature excursion remained below 1°C (peak value of average waveform: 0.4°C; maximum peak value for any pulse: 0.8°C) (Fig. 6(b)), confirming that brief, high-intensity pulses could be reliably delivered with minimal thermal effects at the device shank. We note that the normalized average temperature waveform is well approximated by a 1-D heat kernel 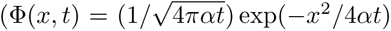, dashed line in Fig. 6(b)) with *α*=4.515 and x=0.285. Looking at the histograms for each pulse in terms of overall power consumption (Fig. 6(c)) and maximum temperature excursion (Fig. 6(d)), we also observed that this level of device heating was representative for all channels, irrespective of the photonic switching tree configuration.

**Fig. 6.**
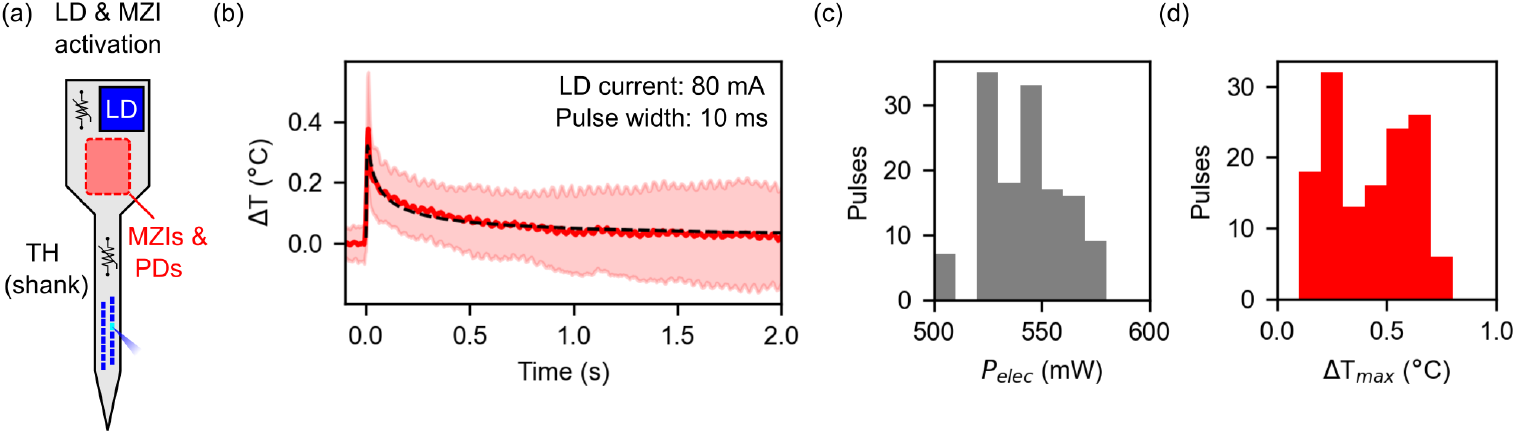
Temperature measurements during extended *in vivo* device operation using integrated thermistors. (a) Conceptual illustration of the neural probe during *in vivo* experiments. Both the LD and photonic switching tree are activated in concert to achieve addressable single-emitter photostimulation. (b) Average relative temperature response (mean ± SD) of the shank thermistor to photostimulation pulses (LD current: 80 mA, pulse width: 10 ms) delivered *in vivo* (n=135 pulses across 16 emitters). Temperature impulse response was modeled using a fitted heat kernel normalized to the maximum temperature excursion (black dashed line). (c) Histogram of combined power dissipation (P*_elec_*) of the LD and photonic switching tree for pulses delivered in (b). (d) Histogram of maximum temperature excursions at the shank thermistor during pulses delivered in (b).

### 2.4 Demonstration of addressable photostimulation and on-chip sensing in head-fixed optogenetic mice

To verify the functionality of the laser-integrated nanophotonic neural probes for optogenetic experiments, we conducted an *in vivo* experiment in an awake, head-fixed optogenetic (Thy1-ChR2^33^) mouse responsive to blue-light photostimulation.

Because the prototype nanophotonic neural probe did not have electrodes for recording electrophysiological activity, we simultaneously implanted a CMOS multielectrode array silicon probe (Neuropixels 1.0) ^29^ in the vicinity of the nanophotonic probe for monitoring photostimulation-evoked responses.

Using two separate micromanipulators, both silicon probes were implanted in cortex within approximately 100 µm of each other (Fig. 7(a)), which was confirmed post-experimentally by imaging the fluorescently-dyed insertion tracks of the probes (Fig. 7(b)). Once implanted, individual photostimulation pulses (amplitude: 40-70 µW, duration: 10 ms, 8-10 pulses per emitter) were delivered sequentially from even-numbered emitters (2-16) while evoked neural activity was monitored using the Neuropixels probe. During photostimulation, we observed highly localized spiking activity across multiple recording channels, confirming that the photostimulation intensity was sufficient for evoking neural activity near the Neuropixels probe (Fig. 7(c)). Following the experiment, we then performed post processing and spike sorting ^34,35^ on the electrophysiological data to assess the responses of individual putative units to targeted photostimulation. After spike sorting, we identified 168 units during the experiment which met our inclusion criteria (see Sec. 4.5). Here, we show 4 units which are representative of the responses observed across all units. Unit 1, which was located near the tip of the Neuropixels probe and away from the nanophotonic probe, had no observable response to photostimulation from any channel (Fig. 7(d)). In contrast, Units 2, 3, and 4, which were located closer to the crossing point between the silicon probes, exhibited localized activation for superficial emitters (Unit 2: emitters 10 and 12; Unit 3: emitters 10, 12, 14, and 16; Unit 4: emitters 10, 12, 14, and 16) but not for deep emitters (Fig. 7(e-g)). We then quantified the firing rate for one of the units (Unit 2) in response to photostimulation from superficial emitters which caused an increase in firing rate (10, 12) and all other emitters (2, 4, 6, 8, 14, 16), and found that the firing rate increased from 7±5 Hz to 138±29 Hz (mean standard error of the mean (SEM), measured 5 ms after pulse onset) for these emitters. (Fig. 7(h)). We also completed a similar experiment in a wild type control mouse (n=1) to test whether spiking was caused by tissue heating, but we did not observe any photostimulation-evoked responses (Fig. S5(a-e)). Overall, these results confirmed that the laser-integrated nanophotonic neural probe could stimulate neural activity with spatial and temporal specificity.

**Fig. 7.**
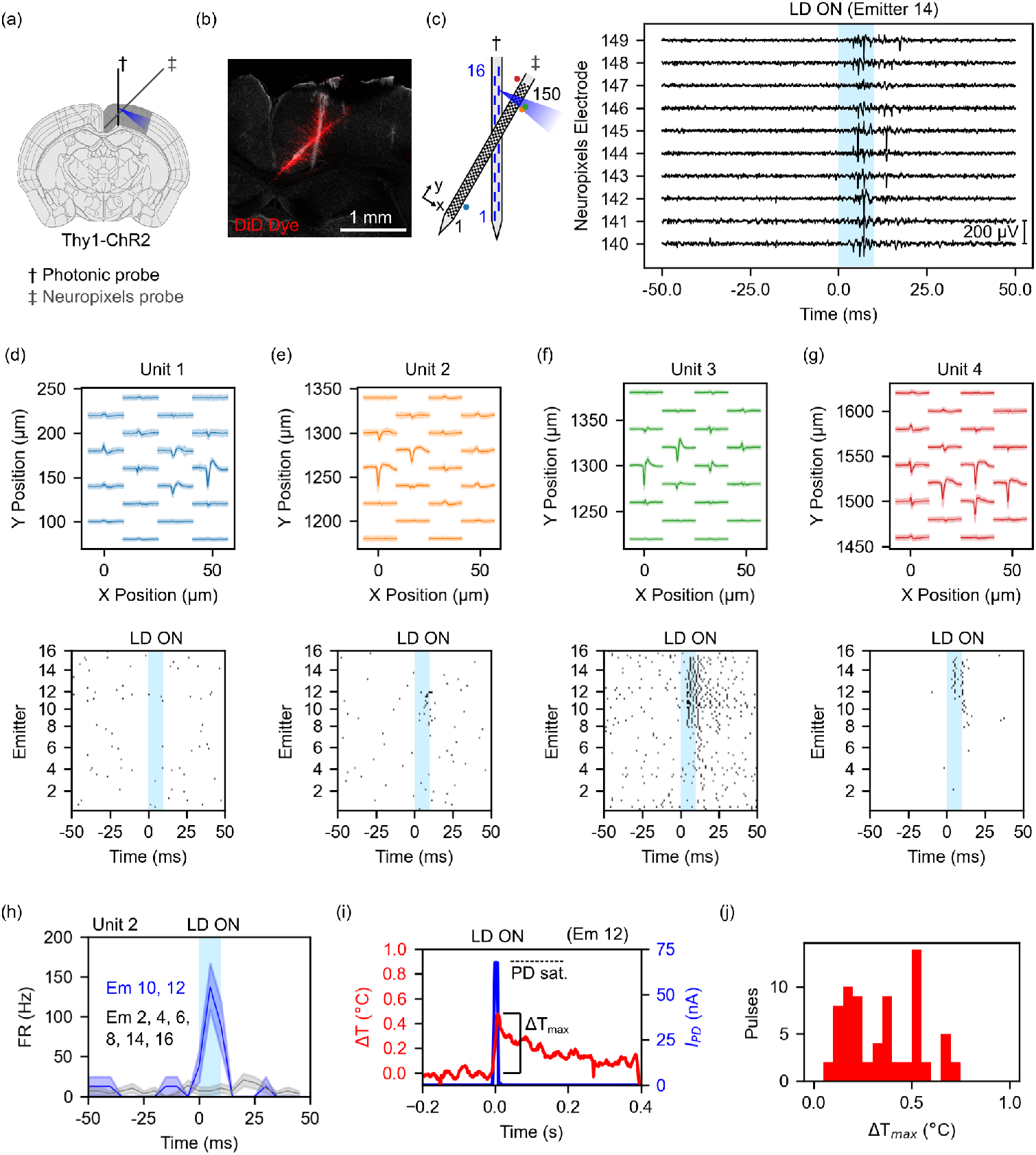
*In vivo* optogenetics experimental demonstration: targeted photostimulation was delivered from the nanophotonic probe and electrical recordings were captured from a Neuropixels 1.0 probe. (a) Conceptual diagram showing the positioning of the two probes. Allen Reference Atlas - Mouse Brain, atlas.brain-map.org ^36^. (b) Histological image showing the fluorescently-dyed insertion tracks of the probes after the experiment. (c) Preprocessed electrophysiological data from the Neuropixels probe during photostimulation. (d-g) Representative putative unit waveforms and spiking activity in response to photostimulation showing (d) no response and (e-g) spatially localized responses. (h) Comparison of firing rate (FR) of Unit 2 in response to photostimulation from specific emitters. (i) Representative photostimulation pulse from emitter 12 with simultaneous PD and shank thermistor measurements. Photocurrent saturation: 67.6 nA (dashed line). (j) Maximum temperature excursions at the shank thermistor for photostimulation pulses (n = 71 pulses from 8 emitters).

Using measurements from the integrated sensors on the nanophotonic neural probe, we then evaluated the device temperature during photostimulation. For each photostimulation pulse, laser onset was marked by a brief activation of the corresponding PD channel and a synchronized increase in the shank temperature (Fig. 7(i)). By analyzing the maximum temperature excursion for all pulses (71 pulses across 8 emitters), we calculated the average maximum temperature excursion as 0.4 °C (Fig. 7(j)), with similar levels of shank heating also observed during the wild type control experiment (Fig. S5(f, g)).

## 3 Discussion and conclusion

In this work, we have developed and validated prototype laser-integrated nanophotonic neural probes for targeted fiberless photostimulation and on-chip sensing, which feature hybrid-integrated InGaN laser diodes, a 16-channel reconfigurable thermo-optic photonic switching tree, and sensors for monitoring on-chip optical power and device temperature. We have also presented our development of a custom peripheral hardware platform for controlling and monitoring these integrated devices, which has been designed to flexibly interface with different neural probe designs, thus streamlining future technology developments. This integrated architecture addresses a key limitation of previous nanophotonic neural probes: the dependence on fiber-coupled external laser and scanning systems, which complicates optical alignment, packaging, operation, and scalability. By combining reconfigurable photonic switching trees with passively aligned flip-chip-bonded LDs in a foundry-fabricated photonic integrated circuit, we transfer key optical power delivery, channel addressing, and monitoring functions from external instrumentation onto the probe itself. In doing so, this work demonstrates a scalable route toward nanophotonic neural probes that integrate the circuit-level functionality required for practical neuroscience experiments while becoming simpler to package, calibrate, monitor, and use.

Through this work, we have shown how integrated PDs can be used to facilitate calibration of the neural probe photonic switching tree and monitor optical outputs during photostimulation. These functionalities are critical for ensuring continuous device operation, as photonic switch bias points may shift due to material aging, necessitating recalibration ^18^, and output powers may drift over time ^17^. A limitation of this work was the range of output powers which could be quantified by the integrated PDs during photostimulation, which was limited by the integration time of the PD ADC (2.2 ms). In principle, the selected ADC (DDC264C) can have a minimum integration time of 320 µs, which could enable output tracking of PD currents up to 469 nA (i.e., emitter powers up to 13.4 µW). With further modifications to the control PCB firmware to increase the clock frequency for this circuit, the system could therefore quantify a wider range of emitter output powers. Alternatively, future implementations could use a different ADC with a greater input current range for tracking greater emitter powers. Lastly, the photonic power taps which couple light into the PDs from the routing waveguides could also be modified to reduce the input light coupling from 5% to below 1% to decrease the PD current and thus enable a wider range of output power monitoring.

We have also shown how integrated thermistors can be used to ensure device temperatures remain within acceptable limits when delivering photostimulation pulse trains. This functionality is especially important for active devices, as neural spiking can be affected by heating of as low as 1 °C ^25,26^. Compared to existing neural probes employing µLEDs, which report maximum heating below 0.6 °C ^2^, our laser-integrated nanophotonic neural probes are able to deliver brief photostimulation pulses with comparable average heating of 0.4 °C at the device shank. Notably, although the electrical power consumed by the LD and photonic switching tree is several orders of magnitude higher than that used by individual µLEDs (500 mW vs. 300 µW), the position of these active circuit elements outside of the brain greatly reduces the effect of heating at the device shank, as we have shown through both simulation (Figs. S3-S4) and experimental measurements (Figs. 5-7). Previous work on simulating laser-integrated neural probes had estimated that temperature excursions caused by laser activation would rapidly increase device temperatures beyond 1 °C and recommended strict management of the laser power and duty cycle to limit device heating ^24^. Here, we have extended this analysis through experimental measurements using integrated sensors and updated simulation models, and found that the temperature at the device shank is less affected by activation of the LD compared to the photonic switching tree for brief optical pulses, owing to the closer proximity of the thermo-optic switches to the device shank and adequate heat sinking of the photonic integrated circuit. For our experiments, we confirm that brief, high-amplitude pulses (emitter output power: 40- 70 µW, pulse duration: 10 ms) can be reliably delivered with minimal heating effects during *in vivo* photostimulation. However, using pulse trains with multiple concurrent photostimulation pulses or longer individual pulses would cause the neural probe shank temperature to rapidly increase. To permit more flexibility in the selection of photostimulation pulse shapes and pulse numbers, future designs may consider changing the probe base shape to increase the surface area for improved heat sinking and move active elements further from the shank (Fig. S4). Likewise, longer implantable shanks may also offer greater thermal resistance compared to shorter shanks, reducing the effects of shank heating from active components on the probe base. Thus, for neural probes using thermo-optic phase shifters for emitter selection, it is crucial to account for device heating caused by the switching tree, and internal temperature sensors act as a critical system component for ensuring device temperatures do not increase by 1 °C. Furthermore, utilizing photonic switches with lower heat generation, such as electro-optic modulators ^37^ or microelectromechanical system (MEMS) switches ^38^, may prove advantageous for enabling continuous device operation with reduced heating, albeit at the expense of more complex fabrication.

In the present design, we have demonstrated on-chip channel selection using a photonic switching tree to address 16 emitters for localized photostimulation. This switching architecture could be scaled further: for example, by adding 1-2 additional layers within the switching tree, this architecture could address 32-64 discrete emitters. Based on the current design, we estimate that these changes would result in (1) a device base area 2-4× larger to accommodate the additional switching layers, (2) an additional 3-6 dB optical loss per switching layer ^20^, and (3) an additional 25-50 mW of electrical power to drive the new thermo-optic switches. Alternatively, to reduce optical losses caused by cascaded switching trees with *>*4 layers, devices could be designed with two or more separate on-chip lasers, each coupled to dedicated 4-layer switching trees, for improved channel addressing with higher emitter optical power. Compared to electrophysiology, where active CMOS-integrated electrodes present the clearest path forward for achieving dense electrode integration (with around 1000 electrodes per shank) ^18,28,29^, achieving a similar density of on-shank emitters remains challenging. Currently, device architectures which support the greatest number of emitters are based on organic light-emitting diodes (OLEDs) (256 emitters per shank) ^3^ and multicore optical fibers (1,200 emitters per fiber) ^11^, yet OLED neural probes are limited to sub-µW optical powers and multicore fiber-based neural probes face barriers to scalability and dense electrophysiology integration. Achieving a similar level of emitter density in nanophotonic neural probes with integrated waveguides would require several innovations in photonic circuit architecture, perhaps through a combination of multiple addressing schemes such as spatial multiplexing ^13,17^, mode multiplexing ^39^, and wavelength-division multiplexing ^14,16,22^.

In conclusion, we have demonstrated foundry-fabricated, laser-integrated nanophotonic neural probes: photonic microsystems with integrated lasers, thermo-optic photonic switches, and on-chip sensors for conducting high spatiotemporal resolution optogenetic experiments, which we have validated through both bench and *in vivo* experiments. By combining both hybrid-integrated LDs and reconfigurable photonic switches, along with integrated sensors for optical power and temperature sensing, our design achieves enhanced scalability in terms of manufacturing, assembly, and circuit complexity. Collectively, this system marks an important step in the development of nanophotonic neurotechnologies, by improving the scalability and usability of these devices to drive advancements in neuroscience research. In future designs, we plan to extend the capabilities of these integrated devices further by increasing the number of emitters for achieving higher-resolution photostimulation, including electrodes for simultaneous electrophysiological recording ^17^, and incorporating multiple laser diodes with distinct wavelengths for facilitating multicolor optogenetic experiments ^16,18^.

## 4 Materials and methods

### 4.1 Integrated nanophotonic neural probe design, fabrication, and packaging

Integrated nanophotonic neural probes were designed and simulated using Lumerical three-dimensional finite-difference time-domain optical simulation (3D FDTD) software (Lumerical, Ansys Inc., Canonsburg, PA, USA) at the Max Planck Institute of Microstructure Physics. Devices were then fabricated at Advanced Micro Foundry (AMF) on 200-mm diameter silicon-on-insulator (SOI) wafers in an active visible-light Si photonics platform. This platform consisted of 120-nm thick silicon nitride (SiN) layer for photonic routing, two aluminum (Al) metal layers with vias for electronic routing, titanium nitride (TiN) resistive heaters for implementing thermo-optic photonic switches, doped Si thermistors for temperature sensing, and waveguide-coupled photodetectors for on-chip optical power monitoring defined in the 220-nm thick SOI layer ^21^. The SiN waveguides were formed using a combination of plasma enhanced chemical vapor depositions and 193-nm deep ultraviolet photolithography. Deep reactive ion etching (DRIE) was used to define the shape of the neural probe chip and devices were thinned by backgrinding to a final thickness of 60 µm. Laser diode (LD) trenches, with mechanical Si stoppers and under-bump metallization, were defined for hybrid integration of flip-chip bonded InGaN lasers ^27,30,31^. LDs were passively aligned and bonded to the neural probes using a sub-micron-precision die bonder (FINEPLACER sigma, Finetech GmbH, Berlin, Germany) with a heated pickup tool (see ref. 30 for additional details). Light was then coupled from the LD to the photonic integrated circuit using an edge coupler with an inverse taper. The junction between the LD and edge coupler was encapsulated with optically-transparent silicone (SYLGARD 184, DOW Inc., Midland, MI, USA) and the LD was then encapsulated in thermally-conductive epoxy (Duralco 128, Cotronics Corp., Brooklyn, NY, USA). The neural probes were mounted on custom-developed carrier PCBs with stainless steel heat sinks using thermally-conductive epoxy (Duralco 132, Cotronics Corp.) and then wirebonded using aluminum wire. Wirebonds were insulated using dielectric epoxy (KATIOBOND GE680, DELO Industrie Klebstoffe GmbH & Co. KGaA, Windach, Germany) and then coated in optically opaque epoxy (EPO-TEK 320, Epoxy Technology, Billerica, MA, United States) to block stray light emission.

### 4.2 Thermistor calibration

Thermistor calibration measurements were completed for a subset of neural probes (n=3) in a heated water bath with respect to a commercially available temperature sensor (accumet AE150, Fisher Scientific, Waltham, MA, USA). Neural probes were inserted in a water bath while the bath temperature was varied using a hotplate. The bath temperature was varied from 20 °C to 45 °C with 5 °C increments while the average temperature of the immersed neural probe shank thermistor was measured. Thermistor temperature measurements were conducted using a 32-bit precision analog-to-digital converter (ADS1263, Texas Instruments) configured for 3-wire resistance temperature device (RTD) measurements. Following calibration measurements, subsequent thermistor measurements were converted to relative temperature values based on the fitted slope of ADC code to temperature values.

### 4.3 Animal surgery and *in vivo* experiments

All animal experiments were conducted under protocols approved by the Animal Care and Use Committee of the University Health Network in accordance with guidelines from the Canadian Council on Animal Care. Adult wild type and transgenic mice (C57BL/6, n=1, age: 2 months, female; Thy1-ChR2-YFP, JAX stock #007612^33^, n=2, age: 2 months, female and male) were used for demonstrating device functionalities *in vivo*. Prior to experiments, mice underwent stereotaxic surgery to implant metal headplates to enable awake, head-fixed recordings. Mice were anesthetized with isoflurane gas (induction: 5%, maintenance: 1-2%) and were injected subcutaneously with buprenorphine (0.1 mg/kg, 0.1 mL) for intraoperative pain management. After reaching the surgical plane of anesthesia, the scalp was cleaned with 70% ethanol and an incision was made along the dorsal surface of the skull. The scalp was then resected, the periosteum was removed, and the exposed skin was pulled away from the incision site and sealed using Vetbond (Fisher Scientific). Stereotaxic coordinates (AP: 0.00 mm, ML: -1.20 mm; AP: 0.00 mm, ML: -2.37 mm) for burr hole craniotomies were marked and mice were then implanted with metal headplates using optically-transparent dental cement (C&B Metabond, Parkell, NY, USA). Following surgery, mice received 1 mL of saline and were allowed to recover for 1-2 weeks prior to experiments.

On the morning of the experiment, mice were anesthetized with isoflurane and received buprenorphine (0.1 mL) for intraoperative pain management. Small, burr hole craniotomies were made on the skull at the marked stereotaxic coordinates to permit two silicon probes (a laser-integrated nanophotonic probe and a Neuropixels 1.0 probe ^29^) to be simultaneously implanted into cortex in the vicinity of motor and somatosensory cortices ^17^) for targeted photostimulation and neural recording. Following the surgery, mice were transferred to a head-fixed experimental setup for awake head-fixed experiments. Prior to implantation, the silicon probes were immersed in DiD fluorescent dye (Vybrant DiD cell-labeling solution, Thermo Fisher) to visualize the insertion tracks after the experiment. Using two micromanipulators (QUAD, Sutter Instruments, Novato, CA, USA; uMp-4, Sensapex, Oulu, Finland), both probes were simultaneously implanted into cortex, with an insertion speed of approximately 200 µm / min. An implantation microscope (InfiniProbe S-25, Infinity Photo-Optical Company, Centennial, CO, USA) and a USB camera were used to guide the positioning and implantation of the two probes. The nanophotonic neural probe was implanted medially (AP: 0.00 mm; ML: -1.20 mm, insertion angle: 90°, depth: 1.3 mm) while the Neuropixels 1.0 probe was implanted laterally (AP: 0.00 mm; ML: -2.37 mm, insertion angle: 48°, depth: 2.2 mm). Photostimulation pulses (amplitude: 10-70 µW, pulse width: 10 ms) were delivered while all PDs and the shank thermistor were monitored.

### 4.4 Histology

Following the experiments, mice were sacrificed and the whole brains were extracted and sectioned coronally with a thickness of 250 µm. Tissue slices were imaged at the Krembil Brain Institute using confocal microscopy (laser excitation wavelength: 637 nm; emission filter: 654 - 815 nm) to visualize the dyed insertion tracks of both probes.

### 4.5 Electrophysiological data processing

Electrophysiological data from the Neuropixels 1.0 probe was preprocessed using SpikeInterface ^34^ and spike sorted using Kilosort4^35^. Preprocessing consisted of a band- pass filter (300 Hz, 6000 Hz), phase shift, and local median subtraction. After spike sorting, units with median amplitudes greater than 50 µV and inter-spike-interval (ISI) violations ratios less than 0.5 were included in the analysis.

## Acknowledgements

This work was supported by the Max Planck Society. D.A.R. was the recipient of a NSERC PGS D award. R.L. and M.J. gratefully acknowledge partial funding support by the German Federal Ministry of Education and Research (BMBF) in the funding program Trusted Electronics (ZEUS) within the joint project VE-Silhouette under grant ME1ZEUS006.

## Contributions

W.D.S. conceived the initial concept for the neural probe technology and designed the integrated neural probe. The fabrication process flow for the neural probes was designed by H.C. and W.D.S. H.C. and G.Q.L. were responsible for the fabrication of the wafers. R.L. and M.J. designed and fabricated the lasers. J.N.S, D.A.R., and W.D.S. conceived, designed, and developed the peripheral electronic system. D.A.R., J.N.S, X.M., F.W., and P.K. developed the probe assembly approach.

D.A.R. conducted all bench experiments. M.S., H.M.C., and D.A.R. prepared and conducted the animal experiments. D.A.R. analyzed all data with feedback from J.K.S.P and W.D.S. D.A.R. wrote the original manuscript with feedback from W.D.S. All authors reviewed and approved the final manuscript. G.Q.L., M.J., J.K.S.P., T.A.V., and W.D.S. supervised the project.

---

**Fig. S1.**
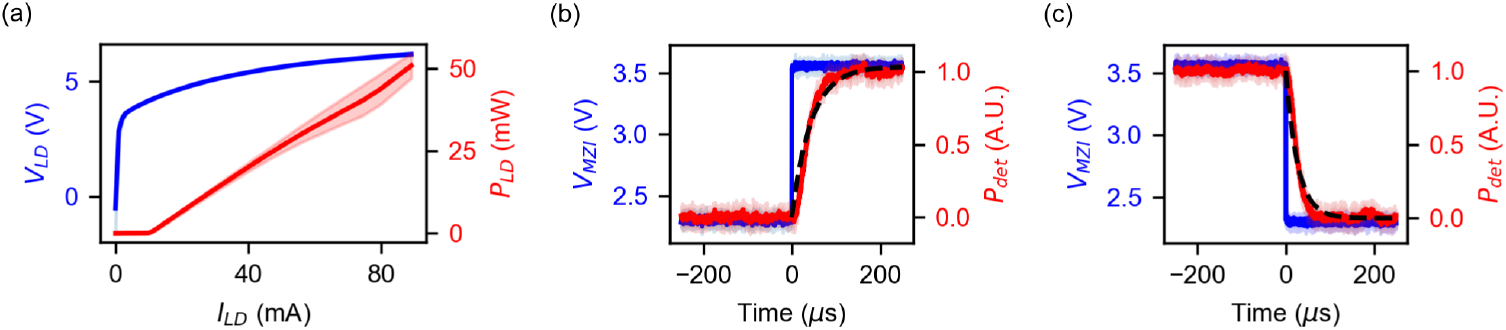
Characterization of laser diodes (LDs) and photonic switches. (a) Light-current-voltage measurements for LD samples on submounts (n=3). (b, c) Rise time (b) and fall time (c) measurements for thermo-optic photonic switch on a neural probe (n=10 measurements from the same photonic switch). Estimated rise and fall times from fitted functions: 96 µs, 59 µs.

**Fig. S2.**
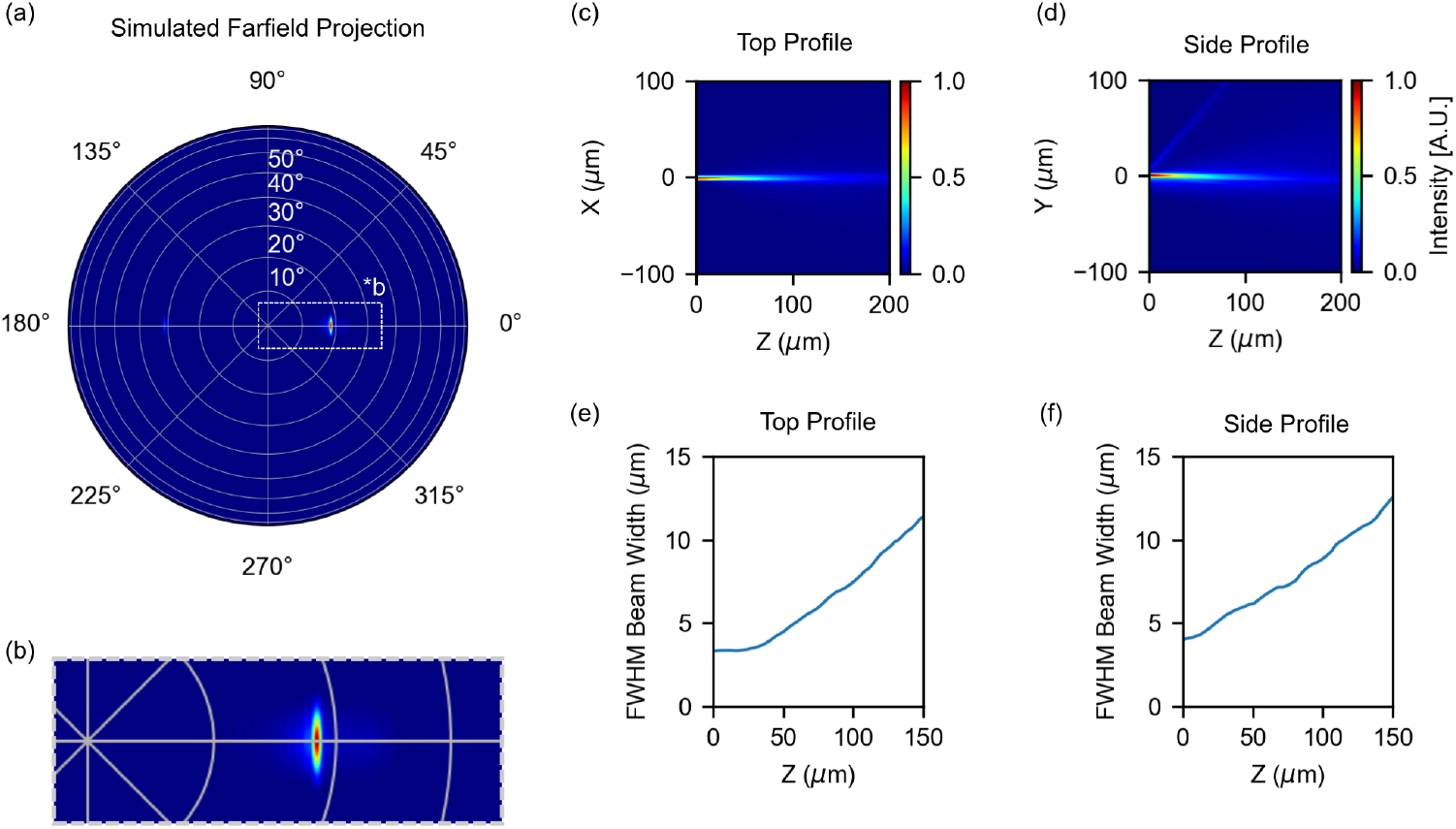
Simulated beam profile of grating coupler emitter in water (n=1.3440). Grating dimensions: width: 6 µm, length: 30 µm, period: 0.4 µm, duty cycle: 50%. (a) Farfield projection of beam profile. (b) Magnified image from (a) showing the primary beam. (c-f) Simulated beam profiles in scattering media (Z: direction of primary beam, X: direction orthogonal to shank axis and grating normal vector, Y: direction orthogonal to X and Z) (c) top profile image, (d) side profile image, (e) full width at half maximum (FWHM) beam width of top profile, (f) FWHM beam width of side profile.

**Fig. S3.**
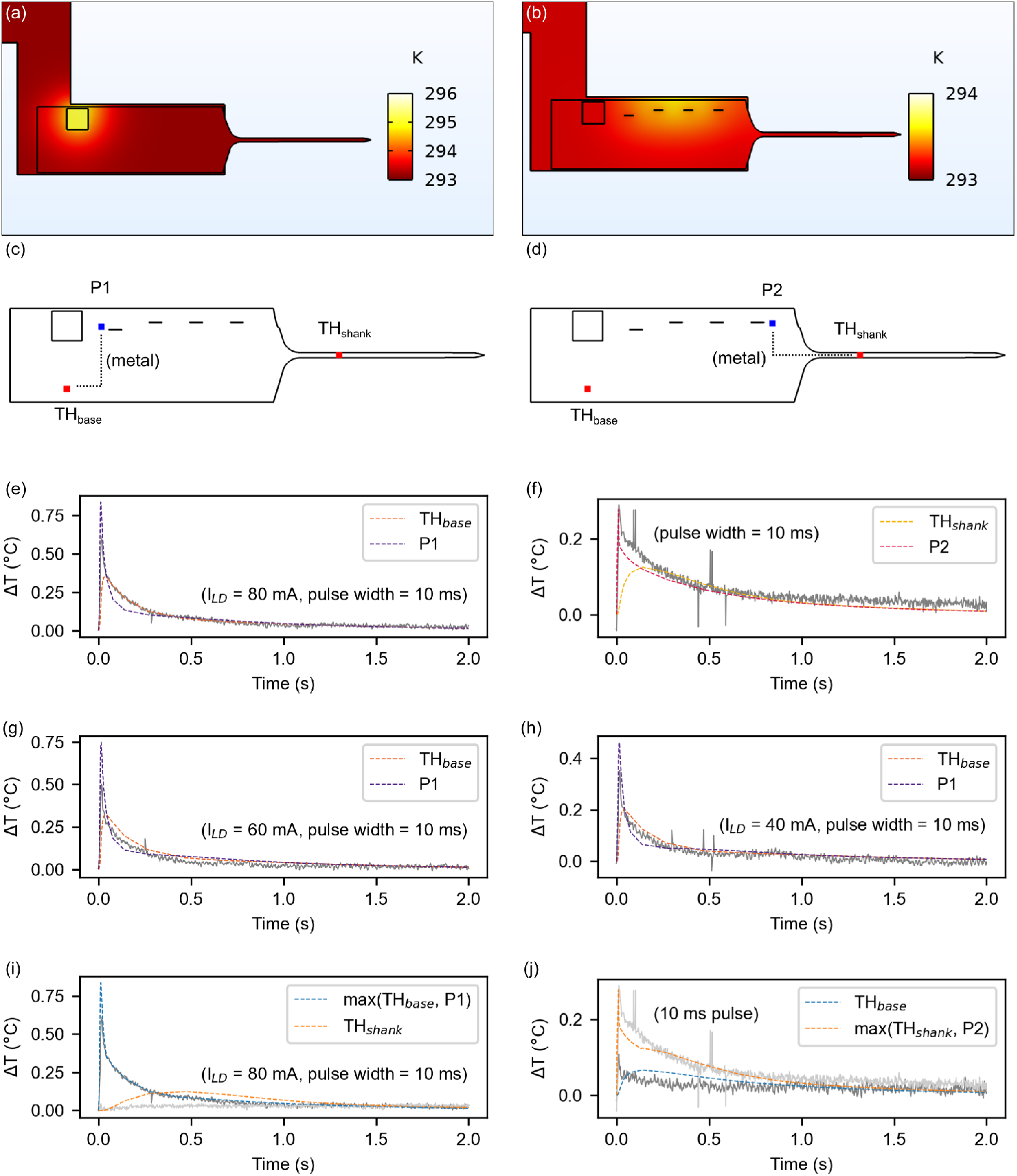
Simulated device heating from isolated laser or photonic switching tree activation. Optimal heat transfer coefficients for the top and bottom surfaces of the heat sink (10*W/*(*m*^2^*K*), 6 ∗ 10^2^*W/*(*m*^2^*K*)) and the top surface of the LD (5 ∗ 10^4^*W/*(*m*^2^*K*)) were identified via parameter sweeps with respect to matched experimental transient data shown in Fig. 5. (a, b) Surface temperature profiles 14 ms after (a) LD pulse or (b) MZI pulse onset. (c, d) Points for measuring thermistor temperatures at device base (*T H_base_*) and shank (*T H_shank_*). Additional points (c) P1 and (d) P2 indicate nearest metal routing which couple heat directly to the thermistors. (e-h) Simulated temperature impulse responses to transient LD activation with currents of (e) 80 mA, (g), 60 mA, and (h) 40 mA and (f) photonic switching tree activation. Matched experimental data from Fig. 5 is plotted in gray. (i, j) Estimated temperature responses at base and shank thermistors for (i) LD and (j) MZI pulses with matched experimental data. Maximum values of (i) *T H_base_*/P1 and (j) *T H_shank_*/P2 are plotted to account for shunt heating through metal routing (metal wiring not directly simulated).

**Fig. S4.**
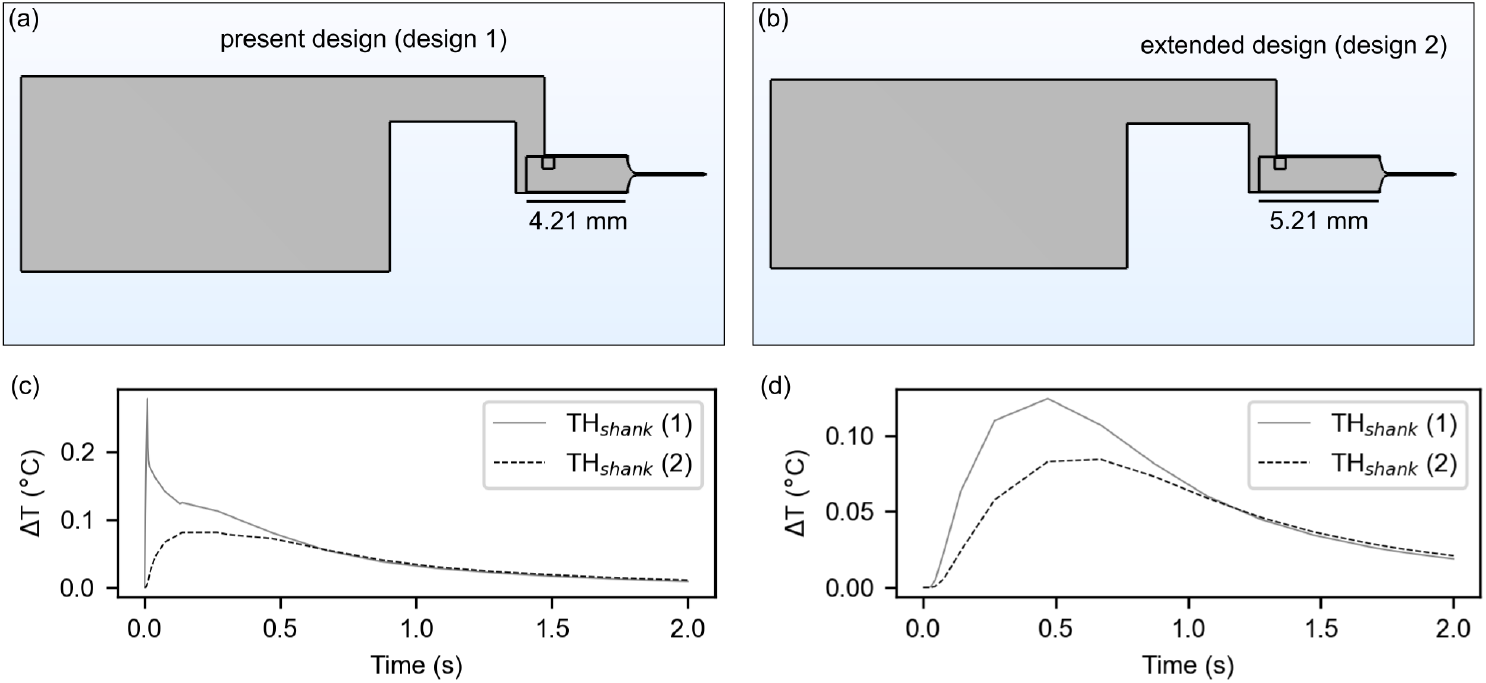
Investigation of device geometry on shank thermistor heating. The present design (a) was compared against a new design (b) with a probe base extended by 1 mm. (c, d) Comparison of shank thermistor heating for designs 1 and 2 in response to pulsed (c) MZI or (d) LD activation. Peak temperature perturbations in response to MZI or LD activation decrease as active elements are positioned further from the shank. Maximum values of *T H_shank_*/P2 are plotted for *T H_shank_* in (c) to account for shunt heating through metal routing (as in Fig. S3(j)).

**Fig. S5.**
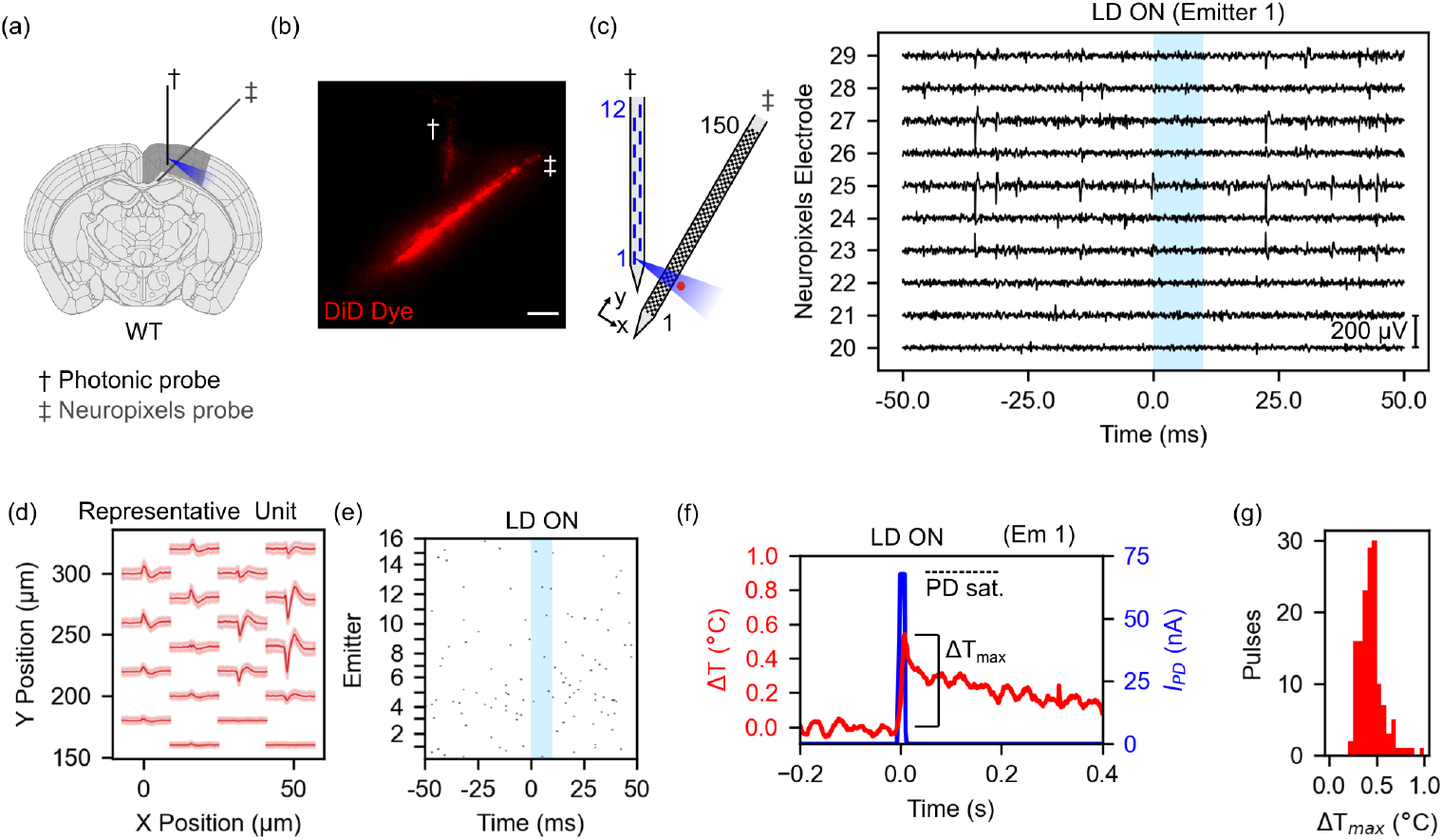
*In vivo* control experiment in a wild type mouse: targeted photostimulation was delivered from the nanophotonic probe and electrical recordings were captured from a Neuropixels 1.0 probe. (a) Conceptual diagram showing the approximate positioning of the two probes. Allen Reference Atlas - Mouse Brain, atlas.brain-map.org ^36^. (b) Histological image showing the fluorescently-dyed insertion tracks of the probes after the experiment. Scale bar: 150 µm. (c) Preprocessed electrophysiological data from the Neuropixels probe during photostimulation. Number of putative units which met the inclusion criteria: n=199 (see Sec. 4.5 for criteria). No photostimulation-evoked responses were observed for any unit. Average emitter output power (measured prior to experiment): 16.7±5.0 µW. Pulses delivered *in vivo*: 8-10 pulses per emitter for all 16 emitters. (d, e) Representative putative unit (d) waveform and (e) spiking activity in response to photostimulation. No response was observed. (f) Representative photostimulation pulse from emitter 1 with simultaneous PD and shank thermistor measurements. Photocurrent saturation: 67.6 nA (dashed line). (g) Maximum temperature excursions at the shank thermistor for photostimulation pulses (mean: 0.4 °C, n = 146 pulses from 16 emitters).

## References

[1] Emiliani, V. et al. Optogenetics for light control of biological systems. Nature Reviews Methods Primers 2, 1–25 (2022).

[2] Vöröslakos, M., et al. HectoSTAR µLED optoelectrodes for large-scale, high-precision in vivo opto-electrophysiology. Advanced Science 9, 2105414 (2022).

[3] Taal, A. J. et al. Optogenetic stimulation probes with single-neuron resolution based on organic LEDs monolithically integrated on CMOS. Nature Electronics 6, 669–679 (2023).

[4] Oh, S. et al. Fiber-less, large-scale opto-electrophysiology interface for micro- scale interaction of multiple brain regions. IEEE Transactions on Biomedical Engineering. 10.1109/TBME.2025.3646326 (2025).

[5] Yan, D. et al. Self-assembled origami neural probes for scalable, multifunctional, three-dimensional neural interface. Preprint at https://www.biorxiv.org/content/ 10.1101/2024.04.25.591141v1 (2024).

[6] Spagnolo, B. et al. Tapered fibertrodes for optoelectrical neural interfacing in small brain volumes with reduced artefacts. Nature Materials 21, 826–835 (2022).

[7] Garwood, I. C. et al. Multifunctional fibers enable modulation of cortical and deep brain activity during cognitive behavior in macaques. Science Advances 9, eadh0974 (2023).

[8] Sahasrabudhe, A. et al. Multifunctional microelectronic fibers enable wireless modulation of gut and brain neural circuits. Nature Biotechnology 42, 892–904 (2024).

[9] Balena, A. et al. Fabrication of nonplanar tapered fibers to integrate optical and electrical signals for neural interfaces in vivo. Nature Protocols 20, 1768–1809 (2025).

[10] Driscoll, N. et al. Multifunctional neural probes enable bidirectional electrical, optical, and chemical recording and stimulation in vivo. Advanced Materials 37, 2408154 (2025).

[11] Yang, S., Yang, K., Chevy, Q., Kepecs, A. & Hu, S. Laser-engineered PRIME fiber for panoramic reconfigurable control of neural activity. Nature Neuroscience 29, 222–233 (2026).

[12] Sacher, W. D. et al. Visible-light silicon nitride waveguide devices and implantable neurophotonic probes on thinned 200 mm silicon wafers. Optics Express 27, 37400–37418 (2019).

[13] Mohanty, A. et al. Reconfigurable nanophotonic silicon probes for sub-millisecond deep-brain optical stimulation. Nature Biomedical Engineering 4, 223–231 (2020).

[14] Lanzio, V. et al. Scalable nanophotonic neural probes for multicolor and on-demand light delivery in brain tissue. Nanotechnology 32, 265201 (2021).

[15] Neutens, P. et al. Dual-wavelength neural probe for simultaneous opto-stimulation and recording fabricated in a monolithically integrated CMOS/photonics technology platform. 2023 International Electron Devices Meeting (IEDM) (2023.

[16] Roszko, D. A. et al. Foundry-fabricated dual-color nanophotonic neural probes for photostimulation and electrophysiological recording. Neurophotonics 12, 025002 (2025).

[17] Chen, F.-D. et al. Implantable nanophotonic neural probes for integrated patterned photostimulation and electrophysiological recording. npj Biosensing 2, 15 (2025).

[18] Lakunina, A. A. et al. Neuropixels Opto: combining high-resolution electrophysiology and optogenetics. Nature Methods 23, 1207–1216 (2026).

[19] Mu, X. et al. Nanophotonic neural probes for in vivo photostimulation, electrophysiology, and microfluidic delivery. Microsystems & Nanoengineering 12, 100 (2026).

[20] Yong, Z. et al. Power-efficient silicon nitride thermo-optic phase shifters for visible light. Optics Express 30, 7225–7237 (2022).

[21] Govdeli, A. et al. Broadband waveguide-coupled photodetectors in a submicrometer-wavelength silicon photonics platform. Optics Express 34, 4709– 4736 (2026).

[22] Segev, E. et al. Patterned photostimulation via visible-wavelength photonic probes for deep brain optogenetics. Neurophotonics 4, 011002 (2016).

[23] Libbrecht, S. et al. Proximal and distal modulation of neural activity by spatially confined optogenetic activation with an integrated high-density optoelectrode. Journal of Neurophysiology 120, 149–161 (2018).

[24] Kampasi, K. et al. Dual color optogenetic control of neural populations using low-noise, multishank optoelectrodes. Microsystems & Nanoengineering 4, 10 (2018).

[25] Stujenske, J. M., Spellman, T. & Gordon, J. A. Modeling the spatiotemporal dynamics of light and heat propagation for in vivo optogenetics. Cell Reports 12, 525–534 (2015).

[26] Owen, S. F., Liu, M. H. & Kreitzer, A. C. Thermal constraints on in vivo optogenetic manipulations. Nature Neuroscience 22, 1061–1065 (2019).

[27] Roszko, D. A. et al. Foundry-fabricated nanophotonic neural probes with integrated lasers, optical switches, photodetectors, and temperature sensors. Proc. SPIE 13836, Optogenetics and Optical Manipulation 2026 138360H (2026).

[28] Yang, X. et al. A CMOS neural probe with 1280 electrodes and 88 emission sites featuring thermo-optic switching and on-chip calibration for dual-wavelength optogenetics. 2026 IEEE International Solid-State Circuits Conference (ISSCC) 616–618 (2026).

[29] Jun, J. J. et al. Fully integrated silicon probes for high-density recording of neural activity. Nature 551, 232–236 (2017).

[30] Mu, X. et al. Hybrid integration of InGaN lasers in a foundry-fabricated visible-light photonics platform. Nanophotonics 15, e70037 (2026).

[31] Nähle, L., et al. Options for cost and performance improvements and integrated emitter design in GaN based electro-optical devices. Proc. SPIE 13386, Light-Emitting Devices, Materials, and Applications *XXIX* 133860C (2025).

[32] Cheng, X. et al. Development of a beam propagation method to simulate the point spread function degradation in scattering media. Optics Letters 44, 4989–4992 (2019).

[33] Arenkiel, B. R. et al. In vivo light-induced activation of neural circuitry in transgenic mice expressing channelrhodopsin-2. Neuron 54, 205–218 (2007).

[34] Buccino, A. P. et al. SpikeInterface, a unified framework for spike sorting. eLife 9, e61834 (2020).

[35] Pachitariu, M., Sridhar, S., Pennington, J. & Stringer, C. Spike sorting with Kilosort4. Nature Methods 21, 914–921 (2024).

[36] Allen Reference Atlas - Mouse Brain [brain atlas]. Available from atlas.brain-map. org.

[37] Liu, T., Bebeti, E., Straguzzi, J. N., Poon, J. K. S. & Sacher, W. D. Visible and near-infrared thin-film lithium niobate modulators with thermo-optic bias. ACS Photonics 10.1021/acsphotonics.6c00668 (2026).

[38] Govdeli, A. et al. Integrated photonic MEMS switch for visible light. Optics Express 33, 650–664 (2025).

[39] Shah, P., Mellouk, E. B., Levine, J. & Mohanty, A. Mode-division multiplexing for visible photonic integrated circuits. Optics Letters 49, 5751–5754 (2024).

